# The Ubiquitin-Proteasome Pathway Mediates Selective Degradation of Unloaded Argonautes in *C. elegans*

**DOI:** 10.1101/2025.09.10.675458

**Authors:** Jenny M. Zhao, Dieu An H. Nguyen, Diego Cervantes, Brandon Vong, Carolyn M. Phillips

## Abstract

Argonaute proteins are essential effectors of small RNA-mediated gene regulation, yet the extent to which their stability depends on small RNA loading remains poorly understood. In *Caenorhabditis elegans*, we systematically disrupted the small RNA binding capacity of multiple Argonaute proteins to assess their stability in the absence of small RNA partners. We found that while most Argonautes remain stable when unable to bind small RNAs, a subset, including PRG-1, HRDE-1, and PPW-2, exhibited markedly reduced protein levels. Focusing on the Piwi-clade Argonaute PRG-1, we show that its destabilization occurs post-translationally and is independent of mRNA expression or translational efficiency. Instead, unbound PRG-1 is targeted for degradation by the ubiquitin-proteasome system. Additionally, the failure to load piRNAs disrupts PRG-1 localization to perinuclear germ granules. We further identify the E3 ubiquitin ligase EEL-1 as a factor promoting degradation of unloaded PRG-1. These findings uncover a critical role for small RNA loading in maintaining the stability and localization of a subset of Argonaute proteins, and reveal a quality control mechanism that selectively eliminates unbound PRG-1 to preserve germline regulatory fidelity.

## Introduction

Precise regulation of gene expression at the transcriptional and post-transcriptional levels is essential for fertility, proper development, and organismal longevity. One fundamental mechanism that contributes to this regulation is RNA interference (RNAi), a conserved biological process in which small RNA molecules recognize complementary target transcripts, resulting in mRNA degradation, translational repression, or transcriptional silencing. In addition to regulating gene expression, RNAi plays critical roles in defending against foreign genetic elements and preserving genome stability (Ghildiyal & Zamore, 2009).

In *Caenorhabditis elegans*, small RNAs are produced via both exogenous and endogenous pathways. Exogenous small interfering RNAs (exo-siRNAs) arise from experimentally introduced or naturally encountered double-stranded RNAs (dsRNAs), a process first described in *C. elegans* by Fire et al. (Fire et al., 1998). These dsRNAs are processed by the RNase III enzyme Dicer into short RNA duplexes (∼21–23 nt), which are then incorporated into RNA-induced silencing complexes (RISCs) (Bernstein et al., 2001; Matranga et al., 2005; Zamore et al., 2000). Endogenous small RNAs, in contrast, are transcribed from genomic loci or generated from mRNA transcripts. These include miRNAs, piRNAs, primary siRNAs, and secondary siRNAs, which are distinguished by their biogenesis pathways, structural features, and associated Argonaute proteins (Ketting & Cochella, 2021).

The core effectors of RNAi are Argonaute proteins, which bind small RNAs to form RISC and mediate gene silencing in a sequence-dependent manner. Argonautes are a highly conserved family of proteins across eukaryotes, characterized by four conserved domains: the N, PAZ, MID, and PIWI domains (Hutvagner & Simard, 2008). The MID and PAZ domains facilitate anchoring of the 5’ and 3’ ends of the small RNA, while the PIWI domain, structurally related to RNase H, can cleave target RNAs when it contains the catalytic DDH triad. In *C. elegans*, however, many Argonautes lack these catalytic residues and instead recruit other silencing machinery to repress gene expression (Yigit et al., 2006).

The *C. elegans* genome encodes 27 Argonaute genes, of which at least 19 are known to produce functional proteins that bind distinct classes of small RNAs (Seroussi et al., 2023; Yigit et al., 2006). These include members of the AGO, Piwi, and WAGO (worm-specific Argonaute) subfamilies, which each have distinct small RNA partners and biological functions. AGO-clade Argonautes include ALG-1, ALG-2 and ALG-5, which primarily bind microRNAs (miRNAs) and RDE-1 that binds primary siRNAs generated from double-stranded RNA and is essential for initiating exogenous RNAi (Brown et al., 2007; Grishok et al., 2001; Tabara et al., 1999). Additional germline-expressed AGO-clade Argonautes, ALG-3, ALG-4, and ERGO-1, bind 26G-RNAs and, along with RDE-1, function as primary Argonautes to trigger the production of 22G secondary siRNAs (Conine et al., 2010; Sijen et al., 2001; Vasale et al., 2010). The single Piwi-clade Argonaute, PRG-1, binds Piwi-interacting RNAs (21U-RNAs) and plays a central role in initiating silencing pathways in the germline (Batista et al., 2008; Das et al., 2008; Wang & Reinke, 2008). WAGO-clade Argonautes act downstream of these pathways by binding 22G-RNAs synthesized by RNA-dependent RNA polymerase (RdRPs) and enforcing gene silencing (Ashe et al., 2012; Luteijn et al., 2012; Shirayama et al., 2012). Cytoplasmic WAGOs (e.g., WAGO-1, PPW-1, PPW-promote post-transcriptional repression, whereas nuclear WAGOs (e.g., HRDE-1 and NRDE-mediate transcriptional silencing and heritable epigenetic regulation (Buckley et al., 2012; W. Gu et al., 2009; Guang et al., 2010). Although structurally related to WAGOs, CSR-1 plays an antagonistic role by licensing gene expression and ensuring proper chromosome segregation (Claycomb et al., 2009; Seth et al., 2013; Wedeles et al., 2013). Together, this diverse Argonaute repertoire enables *C. elegans* to carry out complex, spatially regulated, and highly specific small RNA–mediated gene regulatory programs.

Among the endogenous small RNA pathways in *C. elegans*, the piRNA pathway is distinct in both its biogenesis and function. piRNAs are ∼21 nucleotides in length and are transcribed as single-stranded precursors by RNA polymerase II from thousands of discrete genomic loci (S. G. Gu et al., 2012; Ruby et al., 2006). This transcription requires a dedicated machinery, including components of the small nuclear RNA-activating protein complex (SNPC) and the piRNA-specific factor PRDE-1 (Choi et al., 2021; Kasper et al., 2014; Weick et al., 2014). The resulting precursors undergo tightly regulated 5′ and 3′ end processing, including 2′-O-methylation at the 3′ terminus, to yield mature piRNAs (Billi et al., 2012; Kamminga et al., 2012; Montgomery et al., 2012; Podvalnaya et al., 2023; Tang et al., 2016). Mature piRNAs are then selectively loaded onto PRG-1, which uses partial sequence complementarity to scan for non-self or unlicensed targets in the germline (Bagijn et al., 2012; Lee et al., 2012; Zhang et al., 2018). While direct target cleavage is not required for piRNA-mediated gene silencing, piRNAs initiate a robust secondary siRNA amplification cascade that generates 22G-RNAs, engaging WAGO Argonautes to reinforce gene silencing (Ashe et al., 2012; Bagijn et al., 2012; Lee et al., 2012; Shirayama et al., 2012). This multi-layered surveillance mechanism safeguards genome integrity and ensures transgenerational regulation of gene expression in the germline.

Numerous studies have highlighted the interdependence between Argonaute protein expression and small RNA stability. In *C. elegans*, small RNAs are frequently depleted when their corresponding Argonaute partner is mutated (Batista et al., 2008; Conine et al., 2010; Grishok et al., 2001; W. Gu et al., 2009). Conversely, the expression of some Argonaute proteins is reduced in the absence of their small RNA partners. For example, PRG-1 protein levels are significantly diminished in mutants lacking piRNAs or key piRNA biogenesis factors (Albuquerque et al., 2014; Weick et al., 2014). However, the precise mechanisms underlying PRG-1 regulation remain unclear, and it is not known whether other *C. elegans* Argonaute proteins are similarly affected by the absence of small RNA binding.

In this study, we investigate whether Argonaute protein stability depends on small RNA loading in *C. elegans*. Using a systematic approach to disrupt small RNA binding across a panel of Argonaute proteins, we identify three, PRG-1, HRDE-1, and PPW-2, that are destabilized in the absence of their small RNA partners, while others remain unaffected. We focus on PRG-1 and demonstrate that its degradation occurs post-translationally via the ubiquitin-proteasome system when piRNA loading is impaired. Loss of piRNAs or mutations in PRG-1’s RNA-binding pocket not only reduce protein levels but also disrupt its localization to perinuclear germ granules. Furthermore, we identify the E3 ubiquitin ligase EEL-1 as a factor that promotes degradation of unbound PRG-1. Together, our findings uncover a post-translational quality control mechanism that ensures PRG-1 stability is tightly coupled to piRNA loading, thereby safeguarding the fidelity of piRNA-mediated gene silencing.

## Results

### PRG-1, HRDE-1, and PPW-2 Protein Levels Depend on Small RNA Loading

Argonaute proteins contain two conserved RNA-binding domains that facilitate interactions with their small RNA partners: the MID domain, which binds the 5′ end of the small RNA, and the PAZ domain, which recognizes the 3′ end. The MID domain includes a binding pocket formed by a conserved tyrosine-lysine-glutamine-lysine (Y-K-Q-K) motif that anchors the 5′ phosphate of the small RNA (Ma et al., 2005). This 5′ binding pocket is highly conserved across species (Fig. 1a). Argonaute proteins in the WAGO clade often substitute the first tyrosine residue with histidine, resulting in an H-K-Q-K motif (Fig 1a). This variation may reflect the difference in 5′ end phosphorylation: primary siRNAs are monophosphorylated, whereas secondary siRNAs are triphosphorylated (Pak & Fire, 2007; Ruby et al., 2006).

**Figure 1.**
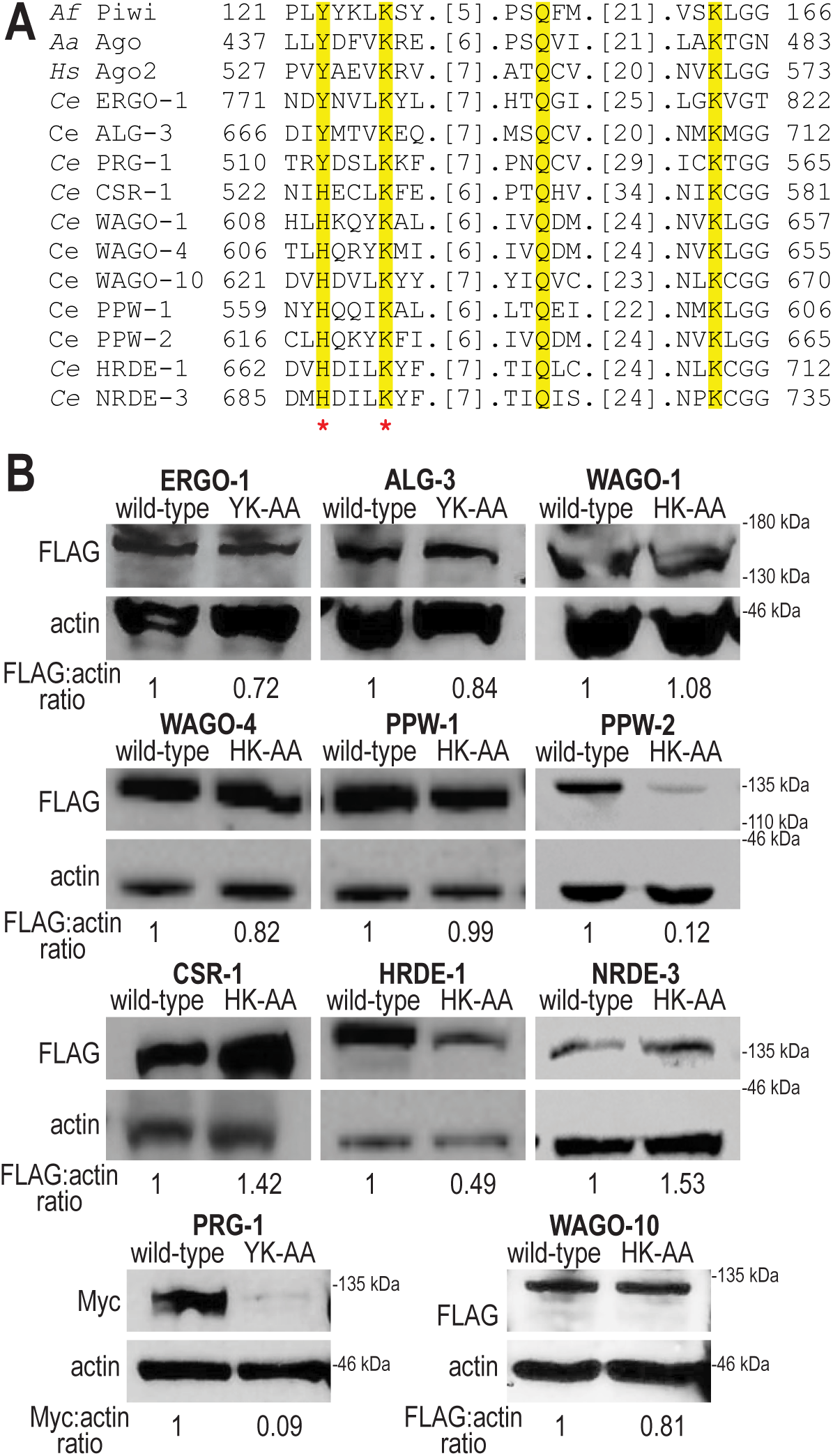
Unloaded PRG-1, HRDE-1, and PPW-2 exhibit reduced expression. **A.** Sequence alignment of the conserved residues (highlighted) in the MID domain of Argonaute proteins across various species. Prefix *Af*, *Archaeoglobus fulgidus*; *Aa*, *Aquifex aeolicus*; *Hs*, *human*; *Ce*, *Caenorhabditis elegans*. Asterisks indicate the two residues that were mutated to alanine to generate small RNA binding-defect Argonaute mutants. **B.** Western blot analysis of wild-type and small RNA binding-defective Argonaute proteins. Anti-FLAG antibodies were used to detect GFP::3xFLAG-tagged ERGO-1, ALG-3, CSR-1, WAGO-1, WAGO-4, PPW-1, PPW-2, HRDE-1, NRDE-3, and 2xFLAG-tagged WAGO-10. Anti-Myc and anti-actin antibodies were used to detect PRG-1 and actin, respectively. Band intensities were quantified using FIJI, and expression levels were calculated as the ratio of Argonaute signal to actin. These ratios were then normalized to wild-type to assess the impact of impaired small RNA binding on Argonaute protein expression.

To investigate the functional consequences of impaired small RNA binding, we engineered small RNA binding-defective mutants representing various Argonaute clades. Previous work showed that mutating the first two residues of the YK/HK motif significantly reduces RNA binding affinity (Ma et al., 2005). Accordingly, we used CRISPR/Cas9 to endogenously tag each Argonaute protein and introduced point mutations replacing the first two residues of the Y/H-K-Q-K motif with alanines (resulting in YK-AA or HK-AA substitutions) (Supplementary Fig. 1a). A semi-quantitative RT-qPCR assay confirmed that the HRDE-1[HK-AA] mutant is defective in small RNA association (Chen & Phillips, 2024). We then performed western blot analysis to compare the expression levels of wild-type Argonaute proteins with their small RNA binding-defective counterparts. Of the eleven Argonautes tested, most, including the primary Argonaute proteins ERGO-1[YK-AA] and ALG-3[YK-AA], as well as secondary WAGO-clade Argonaute proteins CSR-1[HK-AA], WAGO-1[HK-AA], WAGO-4[HK-AA], WAGO-10[HK-AA], PPW-1[HK-AA], PPW-2[HK-AA], and NRDE-3[HK-AA], maintained protein levels comparable to wild-type. In contrast, PRG-1[YK-AA], HRDE-1[HK-AA], and PPW-2[HK-AA] exhibited significantly reduced protein levels: 5–10-fold for PRG-1[YK-AA], 6–8-fold for PPW-2[HK-AA], and 2–4-fold for HRDE-1[HK-AA], compared to their wild-type counterparts (Fig. 1b). The reduction in PRG-1 protein levels observed here is consistent with previous findings showing decreased PRG-1 expression upon loss of piRNA biogenesis factors such as PRDE-1 and PID-1 (Albuquerque et al., 2014; Weick et al., 2014). Together, our results suggest that, while many *C. elegans* Argonaute proteins are stably expressed in the absence of small RNA association, PRG-1, HRDE-1, and PPW-2 require proper small RNA loading to maintain their protein levels.

### piRNA Biogenesis Mutants Recapitulate PRG-1[YK-AA] Destabilization Phenotype

Among the three Argonaute proteins with reduced expression upon failure to load small RNAs, we initially focused on PRG-1, which is a primary Argonaute that specifically binds piRNAs. piRNA biogenesis is a complex process involving both transcription of piRNA loci and processing of precursors into mature piRNAs. These include type I piRNAs, which contain a conserved Ruby motif, and type II piRNAs, which lack this motif (Ruby et al., 2006). Two key biogenesis factors involved in this process are PRDE-1, a nuclear protein required for the transcription of Ruby motif-containing loci and production of type I piRNAs (Weick et al., 2014), and PID-1, which functions more broadly in in global piRNA biogenesis and piRNA precursor accumulation (Albuquerque et al., 2014; Cordeiro Rodrigues et al., 2019).

To determine whether PRG-1 expression is reduced to a similar extent in small RNA binding-defective mutants compared to piRNA biogenesis mutants, we introduced *prde-1(mj207)*, *pid-1(xf35)*, or the *prde-1(mj207); pid-1(xf35)* double mutant into a strain expressing epitope-tagged wild-type PRG-1 and examined protein expression by western blot. To minimize potential effects of larger fluorescent tags on protein folding and stability, we used CRISPR to insert a 2xHA tag at the endogenous *prg-1* locus. Notably, 2xHA::PRG-1[YK-AA] exhibited a dramatic reduction in protein levels, mirroring the results observed with mKate2::3xMyc::PRG-1 (Fig 2a), indicating that the observed reduction in protein levels is not tag-specific. In the *prde-1* mutant background, PRG-1 expression was reduced compared to wild-type but not as severely as in the small RNA binding-defective mutant (approximately 2-fold reduction in *prde-1* vs. 5-fold in PRG-1[YK-AA]). This result likely reflects the fact that *prde-1* is required only for type I piRNA biogenesis, allowing PRG-1 to still bind type II piRNAs. In both the *pid-1* single mutant and the *prde-1; pid-1* double mutant backgrounds, PRG-1 levels were further reduced compared to *prde-1* alone, though still not as depleted as in the PRG-1[YK-AA] mutant (approximately 2.5-fold reduction in *pid-1* single and *prde-1;pid-1* double mutants) (Fig. 2a). These results are consistent with prior findings that low levels of piRNAs are still produced in *pid-1* mutants (Albuquerque et al., 2014), and suggest that residual piRNA association may partially stabilize PRG-1. Overall, these results support the conclusion that PRG-1 levels are reduced when it fails to load piRNAs, and indicates that in the absence of piRNAs, PRG-1 does not efficiently associate with other small RNA classes to maintain its stability.

**Figure 2.**
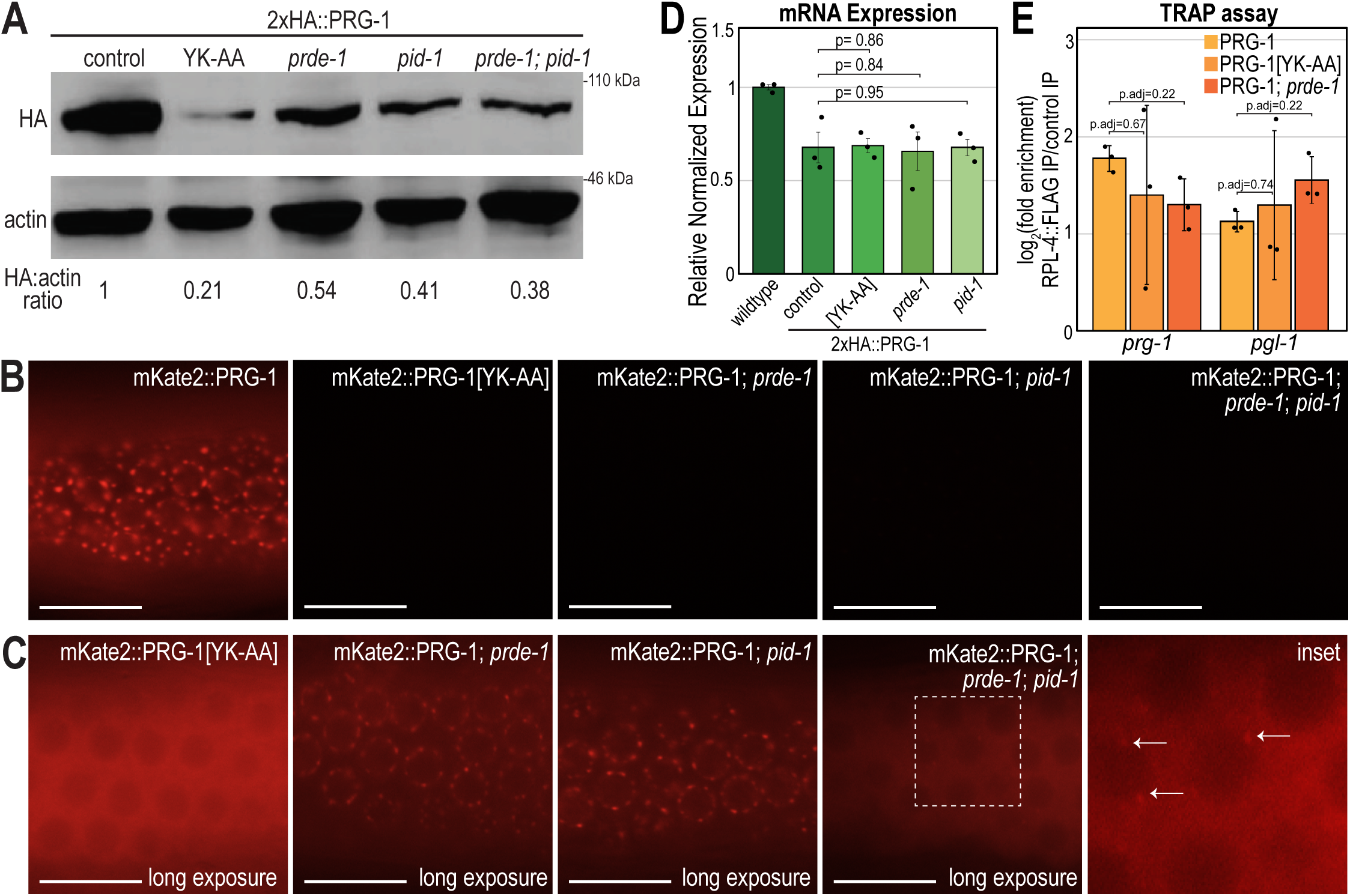
Unloaded PRG-1 is mislocalized and depleted post-translationally. **A.** Western blot showing expression levels of wild-type PRG-1, binding-defective PRG-1 (YK-AA), and PRG-1 in *prde-1(mj207)*, *pid-1(xf35)*, and *prde-1(mj207); pid-1(xf35)* double mutant backgrounds. Actin is shown as a loading control. PRG-1 and actin were detected using anti-HA and anti-actin antibodies, respectively. Band intensities were quantified using FIJI, and expression levels were calculated as the ratio of PRG-1 signal to actin. These ratios were then normalized to wild-type to determine the effect of piRNA binding and biogenesis mutants on PRG-1 protein expression. **B-C.** Live imaging of germlines from animals expressing wild-type PRG-1, binding-defective PRG-1 (YK-AA), or PRG-1 in *prde-1(mj207)*, *pid-1(xf35)*, and *prde-1(mj207); pid-1(xf35)* backgrounds. **B.** Images acquired at identical exposure settings to compare fluorescence intensity. **C.** Images acquired with individually optimized exposures to visualize subcellular localization of residual PRG-1 protein. Scale bars, 15µm. **D.** RT-qPCR of prg-1 transcript levels in untagged wild-type (N2), 2xHA::PRG-1, binding-defective 2xHA::PRG-1[YK-AA], and 2xHA::PRG-1 in *prde-1(mj207)* and *pid-1(xf35)* mutant backgrounds. Expression is normalized to *rpl-32* and relative to expression in wild-type animals. Bar graphs represent the mean of three biological replicates. Error bars indicate standard deviation and *p*-values were calculated using a two-tailed t-test. **E.** RT-qPCR analysis of *prg-1* and *pgl-1* mRNAs in translating ribosome fractions following immunoprecipitation of FLAG-tagged RPL-4 from 2xHA::PRG-1, 2xHA::PRG-1[YK-AA], and 2xHA::PRG-1 in the *prde-1(mj207)* mutant background. Enrichment is normalized to 2xHA::PRG-1 lacking FLAG-tagged RPL-4. Bar graphs represent the mean of three biological replicates, excluding technical replicates where the PCR failed. Error bars indicate standard deviation and *p*-values were calculated using a two-tailed t-test and adjusted for multiple comparisons.

### Altered Localization of Unloaded Argonaute Proteins

We next investigated whether Argonaute protein localization is altered when the proteins are unable to bind small RNAs. Previous studies from our lab and others (Chen & Phillips, 2024; Chen & Phillips, 2025; Guang et al., 2008) have shown that the nuclear Argonautes HRDE-1 and NRDE-3 mislocalize to the cytoplasm when unloaded. Specifically, unloaded HRDE-1 accumulates in the cytoplasm and germ granules, while unloaded NRDE-3 is enriched in the cytoplasm and, specifically during mid-embryogenesis, cytoplasmic foci in somatic cells.

To extend these findings to additional Argonaute proteins, we examined the localization of a subset of Argonautes whose protein levels remained stable in the YK/HK-to-AA mutants. We found that most Argonautes with germ granule localization in the wild-type lost this association when unloaded. For example, both WAGO-1[HK-AA] and ALG-3[YK-AA] failed to localize to germ granules (Supplementary Fig. 1b). In contrast, CSR-1[HK-AA] remained enriched in germ granules, though with increased cytoplasmic localization compared to wild-type CSR-1. Additionally, germ granules in the CSR-1[HK-AA] mutant appeared enlarged and misshapen— phenotypes previously associated with disruption of the CSR-1 pathway (Claycomb et al., 2009; Updike & Strome, 2009; Vought et al., 2005). Notably, ERGO-1 was the only Argonaute examined for which no localization defects were observed in the small RNA binding-defective mutant; both wild-type and YK-AA mutant ERGO-1 remained fully cytoplasmic. Together, these results indicate that proper localization of Argonaute proteins—particularly those that are nuclear or germ granule-associated—depends on their ability to bind small RNAs.

We next examined the localization of PRG-1, which normally localizes to germ granules in the germline (Batista et al., 2008; Wang & Reinke, 2008) (Fig. 2b). In contrast, imaging of the small RNA binding-defective PRG-1 mutant and the piRNA biogenesis mutants under identical exposure settings revealed no detectable PRG-1 signal, further confirming the substantial reduction in PRG-1 protein levels when it is unloaded. To assess the localization of the remaining PRG-1 protein, we increased exposure levels (Fig. 2c). The PRG-1 [YK-AA] mutant showed the most striking change, exhibiting complete loss of germ granule localization and instead displaying diffuse cytoplasmic fluorescence throughout the germline, similar to the mislocalization observed for unloaded WAGO-1 and ALG-3. In the *prde-1* and *pid-1* single mutant backgrounds, PRG-1 retained partial germ granule localization, accompanied by increased diffuse cytoplasmic signal. In the *prde-1; pid-1* double mutant background, PRG-1 localization resembled that of the PRG-1[YK-AA] mutant, with predominantly diffuse cytoplasmic distribution and only occasional perinuclear foci (Fig. 2c). These observations demonstrate that the germ granule localization of PRG-1 depends on its ability to bind small RNAs, supporting the conclusion that small RNA loading is required for proper PRG-1 localization.

### Unloaded PRG-1 levels are reduced post-translationally

To determine whether the instability of unloaded PRG-1 is due to degradation of the *prg-1* mRNA, we first performed qPCR on the small RNA binding-defective mutant and the piRNA biogenesis mutants. Compared to untagged wild-type, both the tagged *prg-1* strain and the mutants showed a modest reduction in *prg-1* transcript levels, suggesting that the endogenous tag may slightly interfere with proper transcription. However, when comparing *prg-1* mRNA levels among the small RNA binding-defective mutant (YK-AA), the piRNA biogenesis mutants (*prde-1* and *pid-1*), and the tagged wild-type strain, we observed no significant differences (Fig. 2d). These results are consistent with previous studies (Albuquerque et al., 2014; Weick et al., 2014) and indicate that the loss of piRNAs or small RNA loading does not significantly impact *prg-1* transcript abundance, supporting a post-transcriptional mechanism of regulation.

Although the reduction of PRG-1 levels appears to occur post-transcriptionally, these results could be due to impaired translation or post-translational degradation. To investigate whether *prg-1(HK-AA)* mRNA is being translated into protein at rates similar to wild-type, we performed a Translating Ribosome Affinity Purification (TRAP) assay. This method, previously shown to capture mRNAs associated with actively translating ribosomes, allows analysis of translation through qRT-PCR of ribosome-bound transcripts (Nousch, 2020). We crossed a FLAG-tagged ribosomal protein (RPL-4) into strains expressing wild-type PRG-1, small RNA binding-defective PRG-1[YK-AA], and PRG-1 in the *prde-1* mutant. RPL-4-associated mRNAs were immunopurified, reverse transcribed, and analyzed by qRT-PCR. As a control, we used *pgl-1*, a P granule marker known to be actively translated in the germline.

We observed enrichment of *pgl-1* mRNA in the TRAP pulldown relative to the negative control (2xHA::PRG-1 without FLAG-RPL-4) across all strains, confirming successful capture of translating ribosomes. Similarly, *prg-1* mRNA was enriched in the pulldown samples from wild-type, PRG-1[YK-AA], and PRG-1 in *prde-1* mutant strains, indicating that *prg-1* is being actively translated in all cases (Fig. 2e). These findings suggest that in the absence of small RNA loading, PRG-1 is synthesized but destabilized at the post-translational level, likely through targeted degradation.

### Degradation of unloaded PRG-1 is not mediated by autophagy

To maintain proteostasis, eukaryotic cells eliminate damaged, misfolded, or excess proteins through two major degradation pathways: the ubiquitin-proteasome system and autophagy. In *Drosophila melanogaster*, a previous study showed that loss of small RNA binding causes Ago1 degradation via the ubiquitin-proteasome system (Smibert et al., 2013), whereas in mice, unbound AGO2 is degraded through autophagy (Martinez & Gregory, 2013). These findings suggest that different Argonaute proteins may be selectively targeted by distinct degradation pathways, and it remains unclear which mechanism is responsible for degrading unbound PRG-1.

To determine the pathway through which unbound PRG-1 is degraded, we inhibited each degradation system, either by RNAi or using pharmacological inhibitors, and assessed PRG-1 protein levels by western blot. If PRG-1 is targeted by either pathway when unloaded, its protein levels should increase following disruption of that pathway.

Autophagy is a multi-step process in which cytoplasmic components are engulfed by autophagosomes, which then fuse with lysosomes to form autolysosomes that degrade the cargo (Klionsky, 2005). To assess whether PRG-1 is degraded via autophagy, we used RNAi to knock down key autophagy components involved at different stages of the pathway: *atg-18*, required for autophagosome formation (Takacs et al., 2019); *epg-5*, a metazoan-specific factor essential for autolysosomal degradation (Tian et al., 2010); and *lgg-2,* which facilitates autophagosome maturation and lysosome tethering (Manil-Ségalen et al., 2014). Western blot analysis revealed that depletion of these factors did not alter PRG-1 protein levels in either the small RNA binding-defective mutant or the piRNA biogenesis mutants (Fig. 3a, Supplementary Fig. 2a), suggesting that autophagy is not responsible for PRG-1 degradation. To further confirm this result, we assessed PRG-1 fluorescence in the germline of *prde-1; pid-1* double mutants following RNAi knockdown of *lgg-1*, an essential autophagy gene involved in autophagosome formation (Manil-Ségalen et al., 2014). As with the western blot results, fluorescence intensity of PRG-1 remained unchanged following *lgg-1* depletion (Fig. 3b,e), indicating that PRG-1 is not degraded via the autophagy pathway when unloaded.

**Figure 3.**
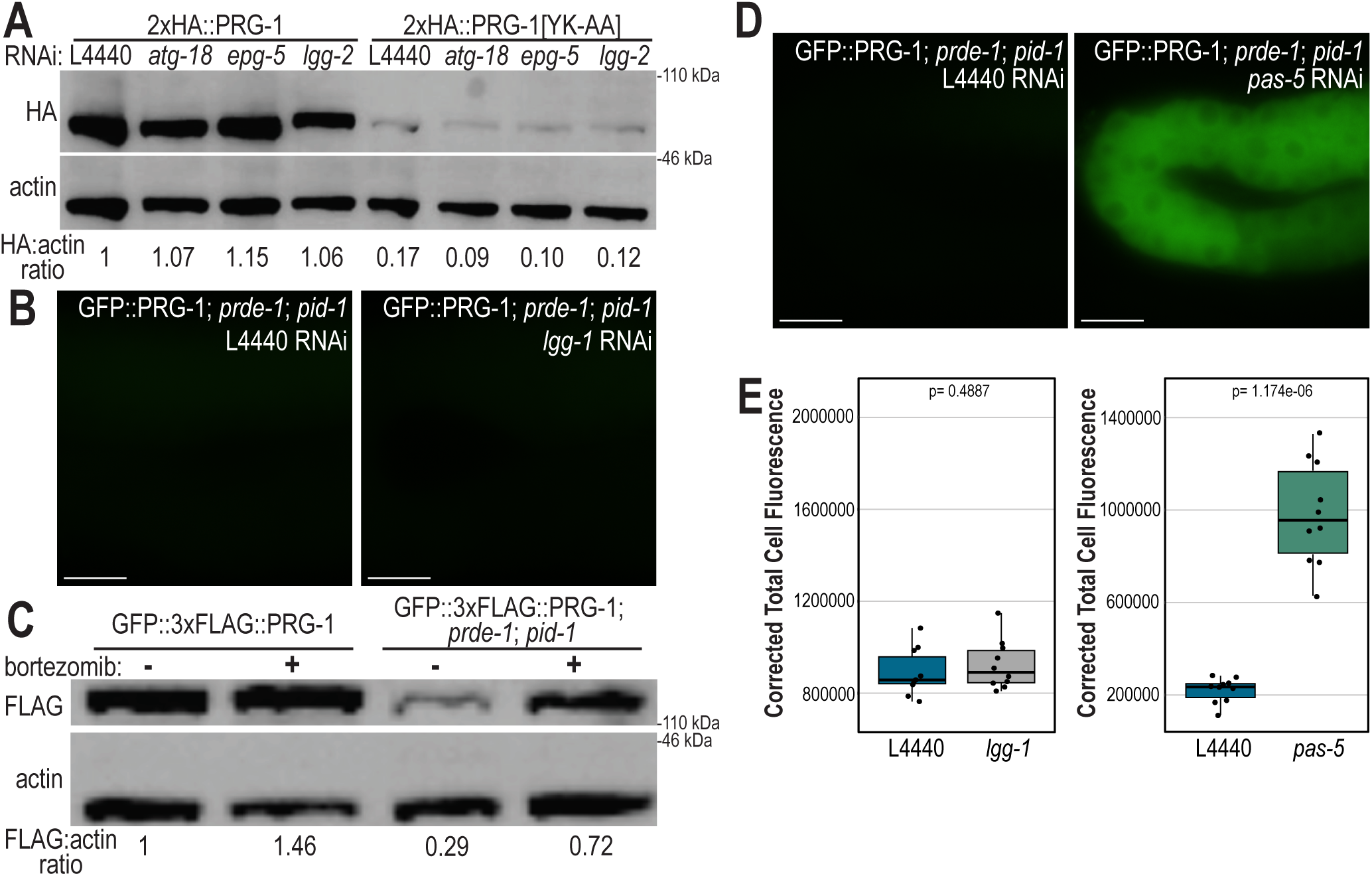
Unloaded PRG-1 is degraded through the ubiquitin-proteasome system. **A.** Western blot of 2xHA::PRG-1 and small RNA binding-defective 2xHA:PRG-1[YK-AA] following RNAi-mediated depletion of autophagy-related genes *atg-18*, *epg-5*, or *lgg-2*. L4440 serves as the RNAi negative control. Actin is shown as a loading control. PRG-1 and actin were detected using anti-HA and anti-actin antibodies, respectively. Band intensities were quantified using FIJI, and expression levels were calculated as the ratio of PRG-1 signal to actin. These ratios were then normalized to wild-type animals on the RNAi negative control (L4440) to determine the effect of knocking down autophagy-related genes on PRG-1 protein expression. **B.** Live imaging of GFP::3xFLAG::PRG-1 in a *prde-1(mj207); pid-1(xf35)* double mutant background following RNAi of either a control (L4440) or the autophagy factor *lgg-1*. Scale bars, 15µm. **C.** Western blot of GFP::3xFLAG::PRG-1 in wild-type and *prde-1(mj207); pid-1(xf35)* double mutant animals treated with 5ug/mL bortezomib to inhibit proteasome activity. Actin is shown as a loading control. PRG-1 and actin were detected using anti-FLAG and anti-actin antibodies, respectively. Band intensities were quantified using FIJI, and expression levels were calculated as the ratio of PRG-1 signal to actin. These ratios were then normalized to untreated wild-type animals to determine the effect of bortezomib on PRG-1 protein expression. **D.** Live imaging of GFP::3xFLAG::PRG-1 in *prde-1(mj207); pid-1(xf35)* double mutant animals following RNAi of a control (L4440) or the proteasome subunit *pas-5*. Scale bars, 15µm. **E.** Quantification of GFP::3xFLAG::PRG-1 fluorescence intensity in *prde-1(mj207); pid-1(xf35)* double mutant animals following RNAi of *lgg-1* (left) or *pas-5* (right). L4440 serves as a negative control. Fluorescence was quantified from 10 regions of interest across 10 individual gonads per condition. Boxes represent the interquartile range (IQR) from the 25th to 75th percentile; whiskers extend to the most extreme data points within 1.5× the IQR. All individual data points are shown. Statistical significance was determined by two-tail t-tests.

### Unloaded PRG-1 is selectively degraded through the ubiquitin-proteasome system

The ubiquitin-proteasome system targets proteins marked for degradation by ubiquitination and breaks them down via the proteasome (Baumeister et al., 1998). To test whether unloaded PRG-1 is subject to proteasomal degradation, we treated animals with bortezomib, a potent proteasome inhibitor (Lehrbach & Ruvkun, 2016). Western blot analysis showed that while wild-type PRG-1 levels are not significantly changed with bortezomib treatment, the levels of unloaded PRG-1 (*prde-1; pid-1* mutant) increased 2.5-fold following proteasome inhibition (Fig. 3c), suggesting that unloaded PRG-1 is indeed degraded through the proteasome. Western blot analysis of HRDE-1[HK-AA] and PPW-2[HK-AA] compared to their wild-type counterparts showed similar results, with levels of unloaded HRDE-1 and PPW-2 increasing 2- and 4-fold, respectively, following proteasome inhibition (Supplementary Fig. 2b), suggesting that unloaded HRDE-1 and unloaded PPW-2 are also degraded through the proteasome.

To corroborate these finding, we examined PRG-1 localization following RNAi depletion of *pas-5*, the *C. elegans* ortholog of a 20S proteasome alpha-type subunit (Kisielnicka et al., 2018; Papaevgeniou & Chondrogianni, 2014). Because *pas-5* knockdown leads to embryonic and larval lethality, RNAi was limited to 48 hours and introduced to worms at the L2 stage. Fluorescence imaging of the germline revealed cytoplasmic accumulation of PRG-1 in the absence of functional proteasome activity (Fig. 3d,e) consistent with impaired degradation. Together, these results demonstrate that in the absence of piRNA loading, PRG-1 is selectively degraded via the ubiquitin-proteasome system rather than autophagy.

### EEL-1 mediates degradation of unloaded PRG-1

Proteins targeted for degradation via the autophagy or ubiquitin-proteasome systems are typically polyubiquitinated, with ubiquitin serving as a molecular tag for recognition and processing. Ubiquitination is catalyzed by a cascade involving three classes of enzymes, with E3 ubiquitin ligases conferring substrate specificity by attaching ubiquitin to target proteins (Hershko & Ciechanover, 1998). To identify factors that may mark unloaded PRG-1 for degradation, we examined previously published PRG-1 immunoprecipitation/mass spectrometry datasets for candidate interacting proteins. From this, we identified 15 genes potentially involved in the ubiquitination pathway (Fig. 4a). To assess their role in PRG-1 regulation, we performed RNAi knockdown of each gene individually in the *prde-1; pid-1* double mutant background and examined PRG-1 fluorescence levels in the germline. Among the 15 candidates, knockdown of *eel-1* using two independent RNAi clones led to a clear increase in GFP::PRG-1 fluorescence, which was confirmed by quantitative analysis (Fig. 4b). *eel-1* encodes an E3 ubiquitin ligase (Ross et al., 2011), suggesting that it may interact with unloaded PRG-1 and facilitate its ubiquitination, thereby targeting it for degradation via the ubiquitin-proteasome system.

**Figure 4.**
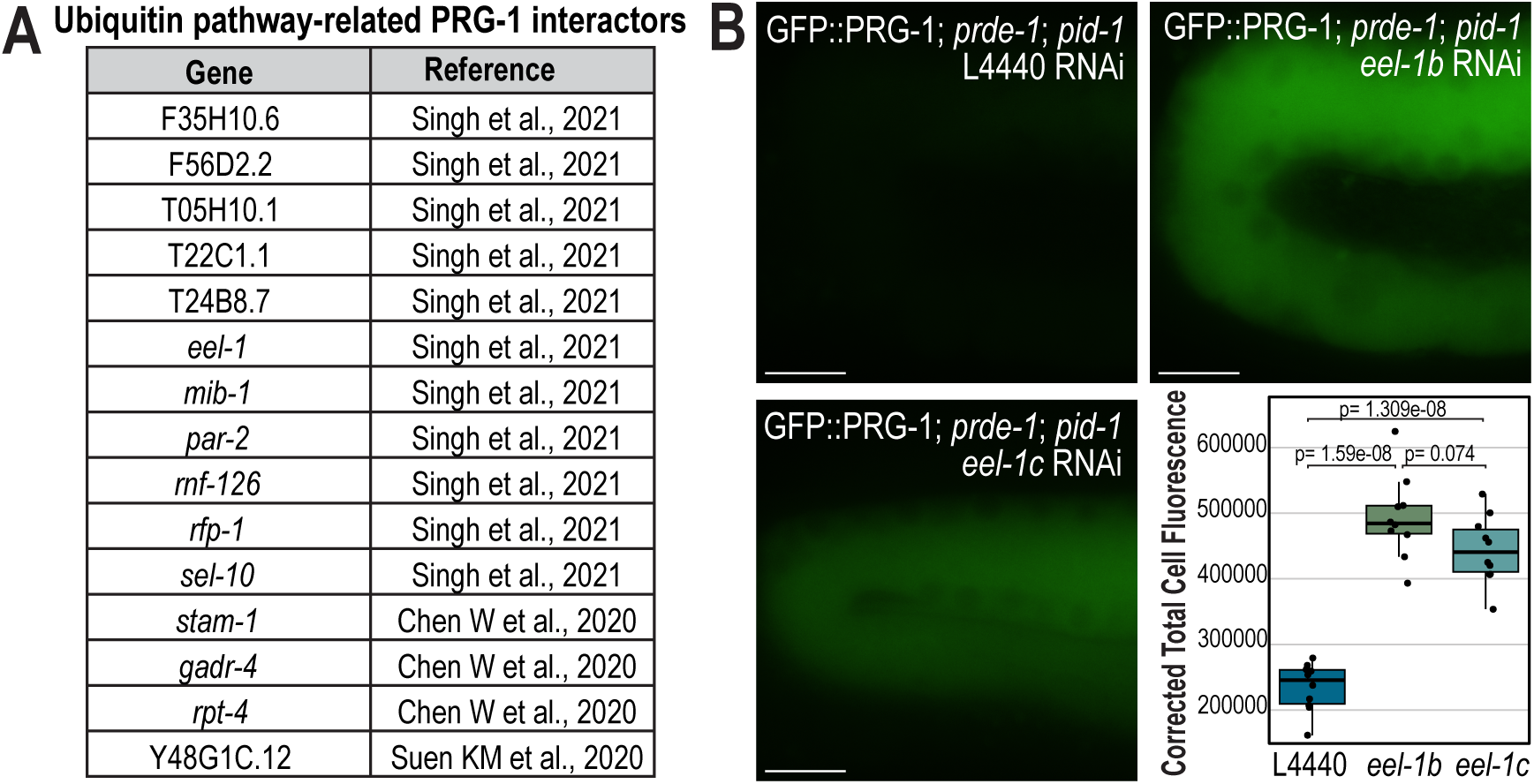
EEL-1 promotes degradation of unloaded PRG-1. **A.** List of candidate *prg-1* interactors involved in the ubiquitin pathway. Genes were compiled from multiple PRG-1 immunoprecipitation–mass spectrometry datasets. **B.** Live imaging of GFP::3xFLAG::PRG-1 in *prde-1(mj207); pid-1(xf35)* double mutant animals following RNAi of a control (L4440) or the E3 ubiquitin ligase *eel-1*. Two RNAi clones targeting distinct exons of *eel-1* (*eel-1b* and *eel-1c*) are shown. Scale bars, 15µm. Quantification of GFP::PRG-1 fluorescence is shown at bottom right. Fluorescence was quantified from 10 regions of interest across 10 individual gonads per condition. Boxes represent the interquartile range (IQR) from the 25th to 75th percentile; whiskers extend to the most extreme data points within 1.5× the IQR. All individual data points are shown. Statistical significance was assessed using two-tail t-tests.

## Discussion

In this study, we identify a subset of *C. elegans* Argonaute proteins, PRG-1, HRDE-1, and PPW-2, that exhibit significantly reduced protein levels when unable to associate with small RNAs. Using site-specific mutations to disrupt small RNA binding, as well as mutants lacking piRNA biogenesis factors, we demonstrate that the expression of these Argonautes is dependent on successful loading with their cognate small RNA partners. Focusing on PRG-1, we find that its reduction in protein levels occurs post-translationally and is not due to differences in mRNA abundance or translation efficiency. Instead, unloaded PRG-1 is selectively degraded by the ubiquitin-proteasome system, as proteasome inhibition restores PRG-1 protein levels. These findings reveal that *C. elegans* employs a quality control mechanism to eliminate unloaded Argonaute proteins, potentially protecting germline function and safeguarding genome integrity.

Previous studies in other organisms have demonstrated that Argonaute proteins are subject to degradation when not loaded with small RNAs (Derrien et al., 2012; Johnston et al., 2010; Kobayashi et al., 2019; Martinez & Gregory, 2013; Smibert et al., 2013; Suzuki et al., 2009). However, the pathways responsible for this degradation vary across Argonautes and species, and the factors that determine which degradation route is engaged remain poorly understood. Here, we identify the E3 ubiquitin ligase EEL-1 as a key player in this process: in *eel-1* mutants, PRG-1 levels are partially restored in the small RNA-binding defective background. These results suggest that EEL-1 acts as a quality control factor, recognizing and promoting the degradation of unloaded PRG-1. Interestingly, *eel-1* knockdown does not fully restore PRG-1 levels, implying that other ubiquitin ligases or degradation mechanisms may act redundantly or in parallel.

In *Drosophila*, a specific lysine residue on Ago1 has been identified as the ubiquitination site in the unloaded state, and this lysine is thought to be buried when Ago1 is bound to small RNAs (Kobayashi et al., 2019), providing an elegant model for selective degradation. Supporting this concept, structural studies suggest that small RNA binding stabilizes Argonaute proteins by bridging flexible domains—particularly between the PAZ and MID domains—thereby masking degradation-prone surfaces (Elkayam et al., 2012). While the specific ubiquitination sites on PRG-1 remain to be identified, it is plausible that, similar to *Drosophila* Ago1, key lysine residues may become exposed only in the unloaded conformation.

In addition to stability, small RNA loading also influences subcellular localization. We find that PRG-1 lacking piRNAs fails to localize to perinuclear germ granules and instead accumulates diffusely throughout the cytoplasm. Similar mislocalization is observed for other Argonautes such as WAGO-1 and ALG-3, which normally localize to germ granules but become cytoplasmic when unloaded. Likewise, nuclear Argonautes HRDE-1 and NRDE-3 are excluded from the nucleus in the absence of small RNAs and relocalize to the cytoplasm (Chen & Phillips, 2024, 2025; Guang et al., 2008). These findings suggest that small RNA loading promotes or stabilizes compartment-specific Argonaute localization, possibly by facilitating interactions with anchoring cofactors or structural components of granules and nuclear transport machinery. However, this coupling is not universal; ERGO-1, for example, retains its cytoplasmic localization in the unloaded state, highlighting pathway-specific differences in localization control.

Collectively, our findings reveal small RNA loading as a key determinant of Argonaute protein fate in *C. elegans*, regulating both their stability and localization. This dependence on small RNA association is enforced by a surveillance mechanism that selectively degrades a subset of unbound Argonautes, thereby ensuring that only properly loaded complexes persist. Such quality control likely serves to maintain small RNA pathway fidelity and protect germline function by preventing the accumulation of non-functional or potentially disruptive Argonaute proteins. Future studies will be needed to identify additional E3 ligases, define the structural basis of degradation signals in the unloaded state, and determine why only a subset of Argonautes is regulated in this manner.

## Materials and Methods

### C. elegans strains

*C. elegans* strains were maintained at 20 °C on NGM plates seeded with OP50 E. coli according to standard conditions unless otherwise stated (Brenner, 1974). All strains used in this project are listed in Supplementary Data 1.

### Plasmid and strain construction

All fluorescent and epitope tags were integrated at the endogenous loci by CRISPR genome editing (Arribere et al., 2014; Dickinson et al., 2013; Friedland et al., 2013; Ward, 2015).

<u>Plasmid-based CRISPR</u>: For all CRISPR insertions of fluorescent tags, we generated homologous repair templates using the primers listed in Supplementary Data 2. *gfp::3xFLAG::ergo-1* was assembled into pDD282 (Addgene #66823) by isothermal assembly according to published protocols (Dickinson et al., 2015; Gibson et al., 2009). To protect the repair template from cleavage, we introduced silent mutations at the site of guide RNA targeting by incorporating these mutations into one of the homology arm primers or, if necessary, by performing site-directed mutagenesis (Dickinson et al., 2013). All guide RNA plasmids were generated by ligating oligos containing the guide RNA sequence into BsaI-digested pRB1017 (Addgene #59936) (Arribere et al., 2014). Guide RNA sequences are provided in Supplementary Data 2. The *gfp::3xFLAG::ergo-1* injection mix included 25-50 ng/μl repair template, 50 ng/μl guide RNA plasmid, 50 ng/μl *eft-3p::cas9-SV40_NLS::tbb-2 3’UTR* (Addgene #46168), and 2.5-10 ng/μl GFP co-injection markers. For *wago-10::2xFLAG* and *2xHA::prg-1* constructs, the injection mix included 50 ng/μl FLAG oligo repair template, 25 ng/μl *wago-10* or *prg-1* guide RNA plasmid, respectively, 20 ng/μl *rol-6* repair template, 25 ng/μl *rol-6* guide RNA plasmid (pJA42, Addgene #59930), and 50 ng/μl *eft-3p::Cas9* (pJW1259, Addgene #61251). All mixes were injected into the wild-type strain. F1 animals with the Rol phenotype were isolated and, for *wago-10::2xFLAG* and *2xHA::prg-1* constructs, genotyped by PCR and restriction digest to identify animals with the correct repair (Arribere et al., 2014). All repair template sequences are provided in Supplementary Data 2.

<u>Protein-based CRISPR:</u> For prg-1(cmp231[YK-AA]), prg-1(cmp301[YK-AA]), alg-3(cmp230[YK-AA]), ergo-1(cmp234[YK-AA]), csr-1(cmp244[HK-AA]), ppw-1(cmp335[HK-AA]), ppw-2(cmp331[HK-AA]), *wago-1(cmp207[HK-AA]), wago-4(cmp332[HK-AA])*, and *wago-10 (cmp240[HK-AA])* mutations, we used an oligo repair template and RNA guide (Supplementary Data 2). All injection mixes included 0.25 μg/μl Cas9 protein (IDT), 100 ng/μl tracrRNA (IDT), 14 ng/μl *dpy-10* crRNA, 42 ng/μl gene-specific crRNA, and 110 ng/μl of each repair template. The *prg-1 [YK-AA]* injection mix was injected into USC1232 (*prg-1(cmp220[(mKate2 +loxP + 3xMyc)::prg-1) I* and USC1273 *(prg-1[cmp227(2xHA::prg-1)] I.* The *alg-3 [YK-AA]* injection mix was injected into USC1092 (*alg-3(cmp155[(GFP + loxP + 3xFLAG)::alg-3]) IV.* The *ergo-1[YK-AA]* injection mix was injected into USC1046 *ergo-1(cmp94[(GFP + loxP +3xFLAG)::ergo-1]) V*. The *csr-1[HK-AA]* injection mix was injected into USC1137 *csr-1(cmp173[(GFP + loxP + 3xFLAG)::csr-1]) IV*. The *ppw-1 [HK-AA]* injection mix was injected into JMC225 *ppw-1(tor119[GFP::3xFLAG::ppw-1c]) I*. The *ppw-2 [HK-AA]* injection mix was injected into JMC221 *ppw-2(tor115[GFP::3xFLAG::ppw-2]) I*. The *wago-1[HK-AA]* injection mix was injected into USC 1362 *wago-1(cmp92[(GFP + loxP + 3xFLAG)::wago-1]) I.* The *wago-4[HK-AA]* injection mix was injected into YY1325 *wago-4(gg620[3xflag::gfp::wago-4]) II*. The *wago-10 [HK-AA]* injection mix was injected into USC1191 (*wago-10(cmp204[wago-10::2xFLAG]) V.* Following injection, F1 animals with the Rol phenotype were isolated and genotyped by PCR to identify heterozygous animals with the mutations of interest, then F2 animals were further singled out to identify homozygous mutant animals. The *csr-1 [HK-AA]* animals are sterile and are maintained in a heterozygous state with the nT1 balancer.

### Imaging and quantification

Live imaging of *C*. *elegans* was performed on 1-day-adult animals (∼72 hours after L1 synchronization) for PRG-1, ERGO-1, WAGO-1, and CSR-1 samples or L4 animals (∼48 hours after L1 synchronization) for ALG-3 samples in either M9 buffer containing sodium azide or immobilized with polystyrene microbeads (PolySciences 00876-15) and 25uM serotonin. Imaging was performed on a DeltaVision Elite microscope (GE Healthcare) using a 60x N.A. 1.42 oil-immersion objective or on a Leica Stellaris 5 confocal microscope using the 63x plan apo N.A 1.40 oil immersion objective. Images were pseudocolored using the Adobe Photoshop.

Quantification of fluorescence intensity was performed in FIJI/ImageJ2 (version 2.14.0). 10 synchronized adult animals were imaged for each sample and fluorescence intensity from the rachis of pachytene region of the germline and for a background region was measured. A standardized region of interest (ROI) was used to compare samples: 127.162 pixels^2^ for L4440 to *lgg-1*, 123.589 pixels^2^ for L4440 to *eel-1b* and *eel-1c*, and 90.006 pixels^2^ for L4440 to *pas-5*. The corrected total cell fluorescence (CTCF) was calculated using the formula CTCF = Integrated Density – (area of ROI x mean background fluorescence).

### Bortezomib treatment

Bortezomib (LC Laboratories B-1408) was solubilized in DMSO at a concentration of 100mg/ml. On the day of bortezomib treatment, plates seeded with OP50 were supplemented with bortezomib to reach a concentration of 5ug/ml. Once plates were dried, synchronized L4 animals were transferred to the supplemented plates and treated for 24 hours prior to harvesting.

### Western blots

*C. elegans* were synchronized at 20°C by bleaching gravid adult animals and maintained as starved L1 larvae for at least 24 hours before plating on OP50. For sample collection, one-day-old adult animals were harvested ∼70 hours at 20 °C after L1 synchronization for PRG-1, ERGO-1, CSR-1, PPW-1, PPW-2, WAGO-1, WAGO-4, HRDE-1, and NRDE-3 samples. L4 animals were harvested 48 hours at 20 °C after L1 synchronization for ALG-3 and WAGO-10 samples. Approximately 400 L4s or 200 gravid adults were loaded per lane. Proteins were resolved on 4-12% Bis-Tris polyacrylamide gels (ThermoFisher NW04122BOX), transferred to nitrocellulose membranes (ThermoFisher LC2001), and probed with mouse anti-Myc 1:1000 (ThermoFisher 13-2500), mouse anti-FLAG 1:1000 (Sigma F1804), rat anti-HA-peroxidase 1:1000 (Roche 12013819001), mouse anti-actin 1:10000 (Abcam ab3280) or mouse anti-actin 1:5000 (ThermoFisher MA5-11869), HRP-labeled anti-mouse IgG Secondary 1:10000 (ThermoFisher A16078) and HRP-labeled anti-rat Secondary 1:10000 (ThermoFisher A18871). For western blot following bortezomib treatment, synchronized L1 larvae were plated on OP50 for 48 hours before being washed off and transferred to plates with OP50 supplemented with bortezomib. Following 24 hours of bortezomib treatment, adult animals were harvested. Western blot quantification was carried out using FIJI/ImageJ2 (version 2.14.0). To isolate the protein bands, images were inverted and the background signal was removed using the Subtract Background function with a rolling ball radius set to 50 pixels. Identical regions of interests (ROIs) were taken for the protein of interests within each blot, and a separate but identical ROI for actin was also used. For each sample, integrated density was measured and normalized to its corresponding actin band. These values were then further normalized relative to the wild-type lane (set to 1) to calculate expression levels.

### Immunoprecipitation of ribosomes

100,000 synchronized adult animals (∼68 h at 20 °C after L1 arrest) were collected in biological triplicates by washing off plates with H_2_O and collected in 0.15M KCl and IP buffer (20mM HEPES pH 7.4, 5mM MgCl_2_, and 15mM KCl) supplemented with 0.5mM DTT, 100ug/mL cycloheximide (Millipore Sigma C7698-1G), 1 protease inhibitor (ThermoFisher A32965), 1% Nonidet P40 substitute, and RNaseOUT (ThermoFisher 10777019). Samples were frozen in liquid nitrogen and homogenized using a mortar and pestle. After further dilution into 0.15M KCl and IP buffer (1:10 packed worms:buffer), insoluble particulate was removed by centrifugation. 10% of sample was taken as “input” and another 10% was used for a BCA assay to determine protein concentration. With the remaining lysate, identical protein concentrations across each sample were used for immunoprecipitation. Immunoprecipitation was performed at 4 °C for 1 h using anti-FLAG Affinity Matrix (Sigma Aldrich A2220), then washed at least three times for 10 minutes in 0.15M KCl and IP buffer. RNA was then isolated from the matrix material as described below.

### RNA isolation and qRT-PCR

For determining mRNA expression levels, synchronized adult animals (∼68 h at 20 °C after L1 arrest) were collected in biological triplicates. RNA was isolated using TRIzol reagent (ThermoFisher 15596018), followed by chloroform extraction and isopropanol precipitation. RNA samples were normalized to 10 μg/μL prior to DNase treatment (TURBO DNA-free kit, ThermoFisher AM1907) and reverse transcribed with SuperScript III Reverse Transcriptase (ThermoFisher 18080-051). All Real-time PCR reactions were performed using the 2x iTaq Universal SYBR Green Supermix (Bio-Rad 1725121), following the manufacturer’s protocols, and run in the CFX96 Touch Real-Time PCR System (Bio-Rad 1855195). Samples were run with three technical replicates for each biological replicate. qPCR to check mRNA levels were normalized to *rpl-32*.

To determine expression levels of mRNAs associated with actively translating ribosomes, RNA was isolated from the matrix material, normalized, and reverse transcribed, as described above. All Real-time PCR reactions were performed using the 2x iTaq Universal SYBR Green Supermix (Bio-Rad 1725121), following the manufacturer’s protocols, and run in the CFX96 Touch Real-Time PCR System (Bio-Rad 1855195). Samples were run with three technical replicates for each biological replicate. Technical replicates with no signal detected or greater than 35 cycles were excluded from analysis. Data was normalized to a control sample lacking the FLAG tag. Primer sequences are available in Supplementary Data 2.

### RNAi assays

For RNAi experiments, control L4440, *lgg-1, lgg-2*, *atg-18*, *epg-5*, *eel-1b*, *eel-1c*, and *pas-5* RNAi *E. coli* clones were sequenced verified and cultured at 37°C for 16 hours. RNAi bacteria were then seeded on fresh RNAi plates. Synchronized L1 animals were transferred to seeded *lgg-1, lgg-2*, *atg-18*, *epg-5*, *eel-1b* and *eel-1c* RNAi plates and raised at 20°C for ∼72hrs. Adult animals on *lgg-2*, *atg-18*, *epg-5* RNAi plates were collected for western blot analysis to assess protein expression levels, and adult animals on *lgg-1, eel-1b* and *eel-1c* were imaged to assess fluorescence intensity changes. For the *pas-5* RNAi assay, synchronized L1 animals were plated on OP50 and raised at 20°C for 24 hours before being washed off the plates and transferred to *pas-5* RNAi plates for 48 hours, and then imaged to assess fluorescence intensity changes.

## Acknowledgements

We thank the members of the Phillips lab for helpful discussions and feedback on the manuscript. Some strains were provided by the CGC, which is funded by NIH Office of Research Infrastructure Programs (P40 OD010440).

## Author contributions

J.W.: Conceptualization, Investigation, Formal analysis, Writing–original draft, Writing–reviewing and editing, Visualization

D.H.N.: Conceptualization, Investigation, Visualization

D.C.: Investigation, Visualization

B.W.: Investigation

C.M.P.: Conceptualization, Formal Analysis, Writing–original draft, Writing–reviewing and editing, Supervision, Funding Acquisition.

